# Improved pathway reconstruction from RNA interference screens by exploiting off-target effects

**DOI:** 10.1101/258319

**Authors:** Sumana Srivatsa, Jack Kuipers, Fabian Schmich, Simone Eicher, Mario Emmenlauer, Christoph Dehio, Niko Beerenwinkel

**Affiliations:** Department of Biosystems Science and Engineering, ETH Zürich, Basel, Switzerland; SIB Swiss Institute of Bioinformatics, Basel, Switzerland; Biozentrum, University of Basel, 4056 Basel, Switzerland

## Abstract

Pathway reconstruction has proven to be an indispensable tool for analyzing the molecular mechanisms of signal transduction underlying cell function. Nested effects models (NEMs) are a class of probabilistic graphical models designed to reconstruct signalling pathways from high-dimensional observations resulting from perturbation experiments, such as RNA interference (RNAi). NEMs assume that the short interfering RNAs (siRNAs) designed to knockdown specific genes are always on-target. However, it has been shown that most siRNAs exhibit strong off-target effects, which further confound the data, resulting in unreliable reconstruction of networks by NEMs. Here, we present an extension of NEMs called probabilistic combinatorial nested effects models (pc-NEMs), which capitalize on the ancillary siRNA off-target effects for network reconstruction from combinatorial gene knockdown data. Our model employs an adaptive simulated annealing search algorithm for simultaneous inference of network structure and error rates inherent to the data. Evaluation of pc-NEMs on simulated data with varying number of phenotypic effects and noise levels demonstrates improved reconstruction compared to classical NEMs. Application to *Bartonella henselae* infection RNAi screening data yielded an eight node network largely in agreement with previous works, and revealed novel binary interactions of direct impact between established components.

**Availability**: The software used for the analysis is freely available as an R package at https://github.com/cbg-ethz/pcNEM.git

## 1 Introduction

Network biology attempts to understand the complex interactions required to maintain the structure and function of a cell through a network-based description. Discerning the relations among different genes provides insight into underlying biological mechanisms and processes. In particular, identifying the defects in different components of signalling pathways involved in diseases can facilitate the discovery of novel drug targets. High-throughput measurements of responses to experimental perturbations of different genes provide a rich set of information to reconstruct the dependencies between them (Molinelli et al., 2013). RNA interference (RNAi) (Fire et al., 1998), CRISPR-Cas9 Knockouts (Shalem et al., 2014), gene knockouts (Hughes et al., 2000), and perturbations from targeted drugs (Molinelli et al., 2013) are some extensively used experimental techniques for observing effects of active perturbations in biological systems. Despite access to high-dimensional phenotypic profiles from such experiments, data-driven inference of intracellular networks remains a key challenge in computational biology.

Several models have been developed to effectively infer signalling networks from perturbation experiments with varying degrees of complexity. A full Bayesian method based on a linear ODE model with a sparsity-enforcing prior on the network was developed by Steinke et al. (2007). Molinelli et al. (2013) proposed an approach to construct context specific, *de novo*, and predictive network models of signalling pathways based on non-linear differential equations from single and paired drug perturbation data. Transitive Reduction and Closure Ensemble (TRaCE), developed by Ud-Dean et al. (2014), generates an ensemble of digraphs from a given gene knockout dataset. The algorithm produces the smallest and the largest networks that bound the complexity of networks in the ensemble, with the edges differing between them deemed uncertain. Franks et al. (2014) developed a compartment specific hierarchical graphical model integrating an ensemble of heterogeneous data sources for refining the network hypothesis.

Another well characterized computational framework for inferring networks from high-dimensional RNAi screens are the nested effects models (NEMs) (Markowetz et al., 2005). NEMs are a class of probabilistic graphical models that aim to infer the connectivity between different perturbed genes using the nested structure of observations. Several extensions of NEMs have been proposed since their conception, including general effects model for inference of DAGs (Tresch et al., 2008), dynamic NEMs for inference from time series data (Fröhlich, Praveen, et al., 2011), NEMix for inference under uncertainty of a stimulation from single-cell observations (Siebourg-Polster et al., 2015), and B-NEMs which combine the use of downstream effects with the higher resolution of signalling pathway structures in Boolean networks (Pirkl et al., 2015), among others.

However, the inference of signalling pathways from perturbation data still remains a challenge. The precision of inference depends on the underlying model assumptions, the computational complexity of the inference algorithm and the availability of relevant information in the data. Often, the information in the data is confounded or unavailable due to several underlying physical, chemical, and biological effects in the experimental approach. One such pervasive effect observed in RNAi screens are the strong off-target effects displayed by the siRNAs (Jackson et al., 2006). Off-target activity of siRNAs involves binding of siRNAs to unintended targets by partial complementarity, leading to unintended silencing and convoluted phenotypes. In general, the estimated cumulative off-target contribution to the observed phenotype is much higher than the on-target contribution (Schmich et al., 2015). Fedorov et al. (2006) showed that off-target effects are capable of inducing strong, and quantifiable toxic phenotypes. Further, they also demonstrated that depending on the specificity of the assay, off-targets can be responsible for as many as 30% of the positives identified in a screen. Thus, in reality the observed effects are a combination of multiple knockdowns, making the interpretation of RNAi screens complex.

In this paper, we propose that taking the ancillary off-target information explicitly into account can improve network inference. Seed sequence dependent siRNA off-target information can be predicted using tools such as TargetScan (Lewis et al., 2005). We developed an extension of NEMs called probabilistic combinatorial nested effects models (pc-NEMs) to infer the underlying network structures using probabilities of combinatorial gene knockdowns (Fig. 1). We demonstrate the power of our approach in a detailed simulation study, and apply our model to *B. henselae* infection screening data.

**Figure 1:**
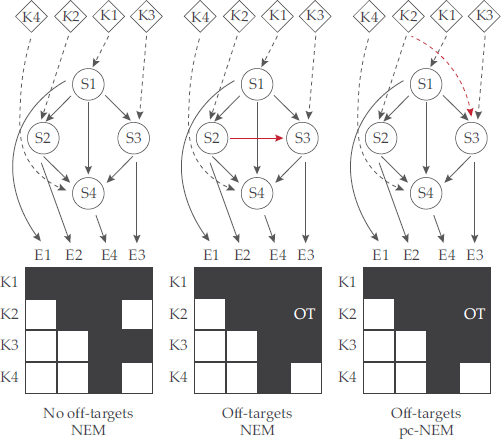
Network inference exploiting off-target effects. A schematic example showing the improvement in inference using off-target information. K1-K4 are the perturbation experiments designed to knockdown S1-S4 genes, and E1-E4 are the observed effects. In the ideal situation with no off-target effects, NEMs successfully infer the true network (left). In the presence of an off-target effect E3 in experiment K2, an additional edge is inferred between S2 and S3 (red solid line), since NEMs assume the perturbations to be strictly on-target (middle). However, by explicitly accounting for the off-target perturbation of S3 in experiment K2 (red dashed line), pc-NEMs can successfully recover the original network (right). The dashed edges are known perturbations and the solid edges are inferred.

## 2 Method

Here, we formally present a novel extension of NEMs called pc-NEMs, and a new inference algorithm based on adaptive simulated annealing. Further, we elaborate on the simulation setup and the analysis of image-based single-cell data for application to infection screening data.

### 2.1 NEM

NEMs are two-layered graph models, designed to infer signalling networks from observations of perturbation experiments. A NEM consists of two types of genes: the perturbed (e.g. silenced) genes called signalling genes (*S*-genes), and the downstream measurable entities called effect genes (*E*-genes) (Markowetz et al., 2005). Let *ε* be a set of *L* effect genes and *S* be a set of *N* signalling genes. Knocking down a specific *S*-gene *S_i_* obstructs the signal flow in the downstream pathway, affecting the *E*-genes attached to *S_i_* and all of its downstream genes. This results in a nested structure of effects which can be used to reconstruct the original signalling graph (Markowetz et al., 2005).

Formally, the dependencies among *S*-genes are given by a binary *N* × *N* adjacency matrix Φ (signalling graph), with Φ*_ij_* = 1 whenever *S*-gene *i* is upstream of *S*-gene *j* for all *i, j* ∊ *S*. The linking of *E*-genes to *S*-genes is formally represented by a *N* × *L* binary matrix Θ (effects graph), with Θ*_se_* = 1 indicating a connection between *E*-gene *e* ∊ *ε* and *S*-gene *s* ∊ *S*. It is assumed that each effect gene is attached to at most one signalling gene, thereby accounting for uninformative effects. Given Φ and Θ, NEMs ascertain that perturbing *S*-gene *s* ∊ *S* leads to an observable downstream effect for *E*-gene *e* ∊ *ε* if there is a path from *s* to *e*, i.e., there exists *s^′^* ∊ *S* such that Φ*_ss_′* = 1, and Θ*_se_* = 1. Thus, mathematically a NEM, *F*, is the product (Tresch et al., 2008),

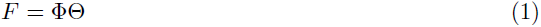

### 2.2 pc-NEM

Off-target effects resulting from non-specific binding of siRNAs, pose one of the biggest challenges in network inference from RNAi screens. We present a novel variation of NEMs called probabilistic combinatorial nested effects models (pc-NEMs), which make use of the combinatorial gene knockdown information resulting from off-targets (Fig. 2). Let *κ* be a set of *K* knockdown experiments. The predicted knockdown information obtained from TargetScan (Lewis et al., 2005) is encoded in a perturbation map *ρ*, which we defined as a *K* × *N* matrix with each entry *ρ_ij_* describing the probability of *S*-gene *j* ∊ *S* being knocked down in experiment *i* ∊ *K*. Given a perturbation map *ρ* and a signalling graph Φ, we then compute the propagation matrix Π, which is also a *K* × *N* matrix with each entry Π*_ks_* describing the probability of *S*-gene *s* ∊ *S* being silenced by experiment *k* ∊ *K*. It is defined recursively as the probability of *S*-gene *s* ∊ *S* or any of its ancestors being silenced in experiment *k* ∊ *K*,

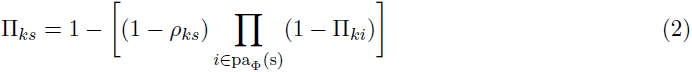

**Figure 2:**
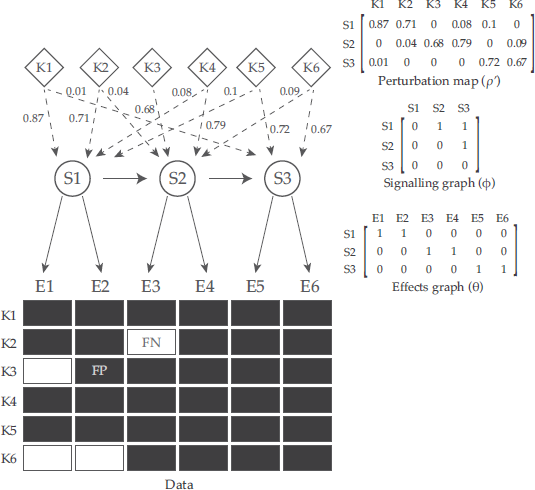
A schematic summary of pc-NEMs. Given the perturbation map *ρ*, a pc-NEM is parameterized by a signalling graph (Φ) encoding the relations between signalling genes (S-genes), and an effects graph (Θ) delineating the attachment of each effects gene (E-gene) to a S-gene. The perturbation map encodes the knockdown probabilities of different S-genes (S1-S3) in each experiment, and is graphically represented by the weighted dashed edges. The data represents the observable effects (E1-E6) for different perturbation experiments (K1-K6), and includes both on-target and off-target observations. By explicitly accounting for the combinatorial perturbation information encoded in *ρ*, pc-NEMs enable successful inference of the signalling graph and the associations between effects and the signalling genes. In addition to off-target effects, the data is subject to noise, resulting in false positive (FP) and false negative (FN) observations, which are modeled using noise parameters α and β, respectively. The dashed edges are known perturbations and the solid edges are to be inferred.

For a given effects graph Θ, the pc-NEM is

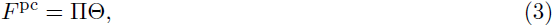

a *K* × *L* matrix with each entry describing the probability of observing an effect in a perturbation experiment. Extending the assumptions of NEMs to pc-NEMs, we require that every *E*-gene can be attached to only one *S*-gene. Thus, each entry 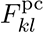 is the probability of observing effect *l* ∊ *ε* in experiment *k* ∊ *K* and is given by Π*_ks_*_(_*_l_*_)_, where the subscript *s*(*l*) denotes that the effect *l* is connected to *S*-gene *s* ∊ *S*.

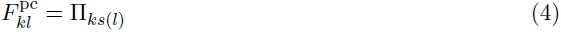

We assume that the observations from perturbation experiments are binary and are stored in a *L* × *K* binary data matrix *D*. Each entry *D_lk_* takes a value of 1 if an effect occurred from the perturbation and 0 otherwise. In an experimental setting, the data is subject to noise. We may observe 0 when there is a true effect from the perturbation with a probability *β* (false negative rate) and 1 when there is no true effect from the perturbation with a probability *α* (false positive rate). For a given perturbation map *ρ*, the probability of each entry *D_lk_* given Φ, Θ, *α*, and *β* is given by

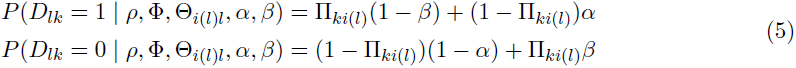

The overall likelihood *P* (*D | F* ^pc^) is then the product of the terms in Eq. 5 across all *L* effects and *K* experiments,

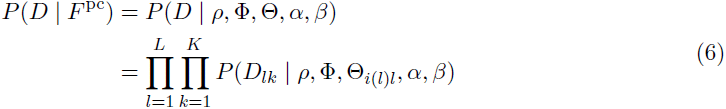

However, as we do not know the topology of the effects graph Θ and we are typically not interested in inferring it from the data, we marginalize the likelihood over all Θ as in the case of classical NEMs (Markowetz et al., 2005). Assuming an uniform prior over topologies, the marginal likelihood is

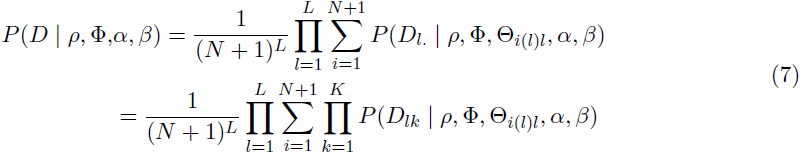

The additional node in the signalling graph accounts for uninformative effects.

### 2.3 Inference algorithm

We developed an inference algorithm maximizing the likelihood (Eq. 7) based on adaptive simulated annealing. In contrast to NEMs, pc-NEMs are not restricted to transitively closed graphs but are defined for all directed acyclic graphs (DAGs). Generally for binary data, the likelihood landscape is very rugged, thus increasing the chances of getting stuck at a local maximum. We employed adaptive simulated annealing (ASA), a global optimization algorithm in which the parameter space can be sampled efficiently, thus making it a good candidate to explore such a rugged likelihood landscape (Ingber, 2000).

ASA is a variant of simulated annealing (SA), which is a metaheuristic for finding global optima with random moves and escaping local optima. The degree of randomness is dictated by the global time-varying temperature parameter *T*. At higher temperatures the system performs more random moves while at lower temperatures the system behaves increasingly analogous to the greedy algorithm. In classical SA, the algorithm begins with a high temperature and is gradually cooled down according to a user defined cooling scheme, ending the run with *T* = 0. In ASA, the temperature parameter is automatically regulated by the algorithm progress, thus making it more flexible and independent of the fixed user defined cooling scheme. The temperature is adapted as

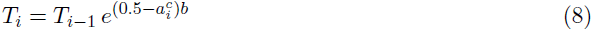

where

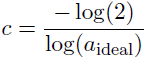

In our implementation of ASA, the temperature is adapted at intervals of 100 steps. The temperature at the end of *i^th^* interval is proportional to the temperature at the end of (*i –* 1)*^th^* interval (*T_i–_*_1_), the acceptance rate for the *i^th^* interval (*a_i_*) and a constant adaptation rate *b* (Eq. 8). The parameter *c* is a scaling constant to ensure a symmetric difference between ideal acceptance rate *a* ideal and *a_i_*. The temperature is adapted based on the location in the landscape and hence is proportional to the acceptance rates. If stuck at a local maximum, the acceptance rates reduce, thereby increasing the temperatures, making the search more randomized. Alternatively, when the acceptance rates are high, the temperatures reduce, simplifying the search to greedy algorithm. The extent of sampling at every step is inversely proportional to the ideal acceptance rate and we chose a default value of 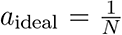. For all the simulation studies, we used default parameters of 20,000 iterations, initial temperature of 50, *a*_ideal_ = 0.125, and adaptation rate *b* = 0.3. Fig. S1 describes the choice of default parameters for the algorithm in detail.

In addition to inferring the network structure, our method can also estimate the error rates *α* and *β* inherent to the data. For inferring both the network and the error rates, the moves stochastically alternate between two distinct spaces, one for the graphs and one for the error rates. For the error rates, the new value is sampled from a bivariate normal distribution with the current value as the mean and a covariance matrix which is updated every 100 steps. For the network structure, the new network is sampled from the neighbourhood of the current DAG, a set of all DAGs generated by adding, deleting and reversing an edge in the current DAG.

The complexity of recursively computing the propagation matrix (Eq. 2) is *𝒪*(*KN*^2^), which is computationally expensive to repeat for every newly sampled graph. However, we can also express the propagation matrix Π, as a function of total number of paths from all the ancestors of a given node. Given a *N* × *N* path count matrix *C* with each entry *C_ij_* denoting the total number of paths (both direct and indirect) from node *i* to node *j*, the total probability of *S*-gene *s* ∊ *S* being silenced by experiment *k* ∊*K* is

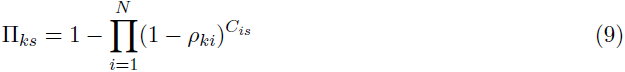

This expression allows us to easily update only the entries affected by the move and hence avoid the recomputation of the entire propagation matrix. Since the moves involve simple operations of addition, deletion or reversal of an edge, at most *N –* 1 nodes are affected and this can be updated at a cost of *𝒪*(*KN*). The model and the inference algorithm were implemented as part of the R/Bioconductor package `nem`.

### 2.4 Data generation for simulation study

We performed simulations to assess the performance of pc-NEMs for varying data set sizes and error rates. TargetScan (Lewis et al., 2005) Version 6.2 was used to predict siRNA off-targets on a genome-wide single-siRNA library from Qiagen. We analysed the predicted siRNA-to-gene target relations matrix with 91,003 siRNAs (rows) and 27,240 genes (columns) (Schmich et al., 2015) to assess the genome-wide off-target distribution. For all simulation studies we used networks sampled from the KEGG pathway database (Kanehisa et al., 2000). In order to exploit the off-target effects for network inference from combinatorial knockdown data, we were interested in sub-networks with the corresponding siRNAs exhibiting a large degree of off-targets. Since the siRNAs targeting the genes involved in different pathways do not exhibit the same degree of off-targets, we first ranked the KEGG pathways in decreasing order of off-target frequencies. For the prioritisation, we first calculated the weight of each gene *i*,

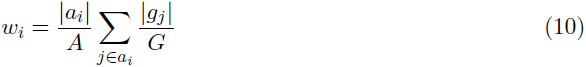

where *A* is the number of siRNAs (91,003) and *G* is the number of genes (27,240) from the target relations matrix, *g_j_* is the set of genes knocked down by siRNA *j* and *a_i_* is the set of siRNAs knocking down gene *i*. The weight of each gene is proportional to the number of experiments perturbing that gene and the degree of off-targets displayed by those experiments. The score for each pathway *P_k_*, was then defined as the difference of weights of genes within and outside the pathway.

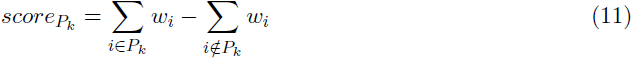

The crux of this scoring function is to weigh heavier off-target dense pathways or pathways with genes that have been perturbed by large number of siRNAs, thus maximizing the combinatorial knockdown within the network. While pathway *hsa*01100 corresponding to the whole network of ”Metabolic pathways” ranked first in this prioritisation scheme, it was not suitable due to its large size. Thus, we chose the second ranked pathway, *hsa*05200, which is titled ”Pathways in cancer”. Further, to evaluate the dependency on the density of off-target effects, we also used the least scoring pathway *hsa*00030, which is titled ”Pentose phosphate pathway”. We sampled 30 sub-networks, each consisting of eight signalling genes from both pathways, using random walks. Subsequently, we derived the corresponding perturbation maps from the target relations matrix for each of these sub-networks. To assess robustness of our model to uncertainty in knockdown strength predictions by TargetScan, we used different perturbation maps for data generation and network inference. We used the perturbation maps derived from target relations matrix for inference and simulated perturbation maps for generating the data. For the simulated perturbation maps, each entry 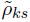 was an independent and identically distributed random variable drawn from a beta distribution with mean equal to the predicted knockdown strength (*ρ_ks_*) and variance inversely proportional to scaling factor *d*, equal to 100,

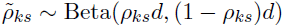

For performance assessment of pc-NEMs in comparison to NEMs, we generated data from transitive closure of the 30 DAGs sampled from *hsa*05200. In order to evaluate the performance as a function of the number of phenotypic effects, we used a fixed, randomly generated effects graph with *L* ∊ {32, 64, 120, 320} for all networks such that the prior probability of attachment of *E*-genes to *S*-genes was uniform. We appended new random phenotypic effects to existing effects, assigning 10% of the effects to uninformative effects. Given the network, simulated perturbation map and effects graph, we simulated binary data. In order to compare the performance as a function of noise, we repeated this process for noise levels (*α*, *β*) ∊ {(0.01, 0.01), (0.20, 0.05), (0.05, 0.20)*}*.

We setup another study for estimating the error rates from the data. We used the same 30 DAGs from *hsa*05200 and for each DAG we sampled four pairs of error rates given by independently and identically distributed random variables drawn from unif{0, 0.5}, to generate 120 different data sets each with 320 phenotypic effects. In the last study evaluating the performance of network inference as a function of noise, for each pair of *α* and *β* in {0.01, 0.1, 0.2, 0.3, 0.4, 0.45, 0.49, 0.5}, we generated datasets with 320 phenotypic effects and the same 30 DAGs from *hsa*05200.

### 2.5 Infection screening data

The data was derived from microscopy image-based infection assays where ATCC HeLa cells were transfected with a genome-wide single-siRNA library from Qiagen followed by infection with *B. henselae*. The cells were then fixed, stained and imaged. The images were corrected for illumination distortion using CIDRE (Smith et al., 2015). Subsequently, the cell features (phenotypic effects) were extracted from the grid of 9 images per knockdown experiment using screeningBee CellProfiler an in-house image analysis solution based on CellProfiler (Carpenter et al., 2006). Features were grouped based on their source segmented objects (parts of the cell), which include: Cells (cell body), Nuclei (cell nuclei), and Perinuclei (perinuclear space). In addition, features derived from Voronoi segmentation of the images were included.

The Qiagen siRNA library typically consists of four different siRNAs per gene, with the exception of talin1 and Cdc42, which had three and eight siRNAs, respectively. We used TargetScan’s (Lewis et al., 2005) predicted off-targets to define the perturbation map for the eight genes across the 35 experiments.

In order to convert single-cell data to gene-level binary data, we first applied B-score normalization to correct for row, column, and plate effects. Then we further normalised the data using MARS (multivariate adaptive regression splines) and z-scoring. This entire process was performed using the R package `singleCellFeatures` (Bennet, 2015). We defined a control distribution because the biological controls were subject to strong edge effects from the experimental setup. The first and last two columns of wells constituted the control wells. We performed Wilcoxon tests between all pairs of control populations and generated a distribution of p-values for each feature, choosing the lower 5*^th^* percentile as the critical p-value. This was done to capture the differences across control populations. For the gene-level data, the knockdown populations were compared to six random control populations (≈ 10% of control wells) using a Wilcoxon test. The resulting p-values were combined using Fischer’s method, and this meta p-value was compared to the critical p-value. The feature was significant and took a value of 1 if the meta p-value was less than the critical p-value and 0 otherwise. The resulting gene-level binary data set consisted of 288 features across 35 knockdown experiments.

## 3 Results

We have developed pc-NEM, a new model based on NEM, which probabilistically models combinatorial perturbations. We first show that pc-NEMs are identifiable and then assess the performance of pc-NEMs in an extensive simulation study. Finally, we apply the model to infection screening data.

### 3.1 Identifiability of pc-NEMs

The classical NEMs reconstruct networks from subset relations or nested data, and are identifiable only in the space of transitively closed networks. Tresch et al. (2008) relaxed the constraint of nestedness in the data and extended the model to general effects models defined over the space of all DAGs. This implied that the perturbation of every parent only affected the parent and its immediate children. However, the interpretation of such reconstructed networks is questionable in the context of biological signalling pathways as signals propagate downstream, beyond the immediate neighbours. Our model overcomes the limitations of NEMs and general effects models by inferring DAGs from nested data. Modelling the perturbations probabilistically makes pc-NEMs identifiable over the entire DAG space. Thus, pc-NEMs can be used to distinguish between equivalent DAG structures from nested data, as opposed to classical NEMs or general effects models. The detailed proof is provided in the appendix.

### 3.2 Performance evaluation on simulated data

Since NEMs can only infer transitively closed networks, we restricted the comparison of the performance of NEMs and pc-NEMs with varying number of effects and noise levels to data generated from transitively closed networks (Fig. 3). We used structural Hamming distance (SHD) between the inferred graph and the KEGG graph as a measure of performance. Each panel in Fig. 3 describes the performance of the models for inferring networks from data with fixed noise levels and increasing number of effects. Since we developed a new model and a novel inference algorithm, we first ran NEMs with the module network algorithm (Fröhlich, Fellmann, et al., 2008), and NEMs with adaptive simulated annealing and compared their performances to performance of pc-NEMs with adaptive simulated annealing. We chose module network as it performed best among all other inbuilt algorithms in the `nem` package (Fig. S2). In general we observed an increase in performance (decrease in SHD) with increasing number of effects across all noise levels. With as few as 64 effects Next, we relaxed the transitive closure constraint and extended the simulation study to the entire DAG space. Although general effects models are not restricted to transitively closed networks (Tresch et al., 2008), as the data generated is still nested it would always infer a transitively closed network, providing pc-NEMs with a huge advantage over them *a priori*. Thus, we assessed the performance of only pc-NEMs and observed that the performance improved with increasing number of effects. In order to study the effect of degree of off-targets on performance, we performed a simulation study for inferring networks from KEGG pathways with high (hsa05200) and low (hsa00030) densities of off-targets (Fig. 4A) and varying number of effects and noise levels. The average numbers of off-targets per network for the high and low density pathways were 28.27 and 9.33, respectively. The performance of pc-NEMs once again improved with increasing number of effects and was independent of the frequency of off-targets at a number of 320 effects. While the variance was smaller for reduced off-target effects, the overall performance was consistent across noise levels and frequencies of off-target effects. In general, the simulation studies show that using the additional information of siRNA off-target effects encoded in the perturbation map improves the network inference from combinatorial gene knockdown data.

**Figure 3:**
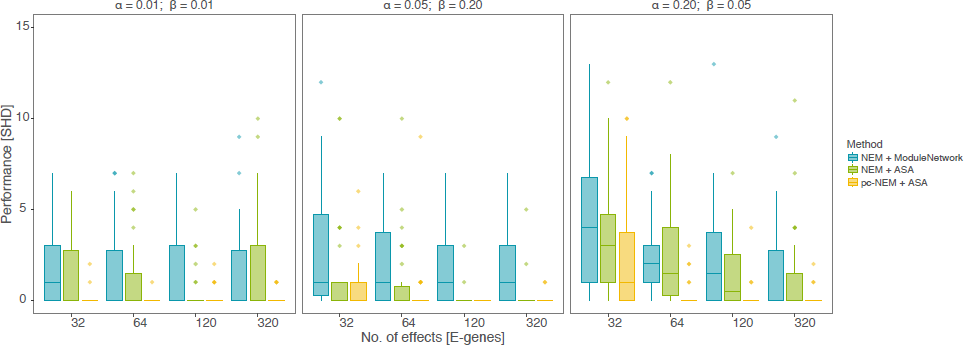
Results from simulation study on transitively closed networks. Structural Hamming distance (SHD) measuring the performance (y-axis) of pc-NEMs (yellow) and NEMs with adaptive simulated annealing (green) and module network (blue) algorithms, on simulated data from 30 different transitively closed networks of size *N* = 8 and perturbation maps, with varying number of phenotypic effects and noise levels.

So far it was assumed in all the simulations that the noise parameters were known, i.e., the networks were inferred with the same noise parameters used for generating the data. Since pc-NEMs can also estimate error rates, we set up a simulation study to assess the performance of inferring the error rates from 120 different pairs of error rates (Fig. 4B). The maximum likelihood estimates of the errors had a Pearson’s correlation of 0.967 and 0.784 for *α* and *β*, respectively, with their true values. The outliers typically correspond to data sets with high values of both *α* and *β*. The general trend of slight overestimation of parameters, especially *β*, is a consequence of imbalanced data sets with more zeroes than ones and reduced accuracy of network inference. However, given the true network, there is no bias in noise parameter estimation (Fig. S3).

**Figure 4:**
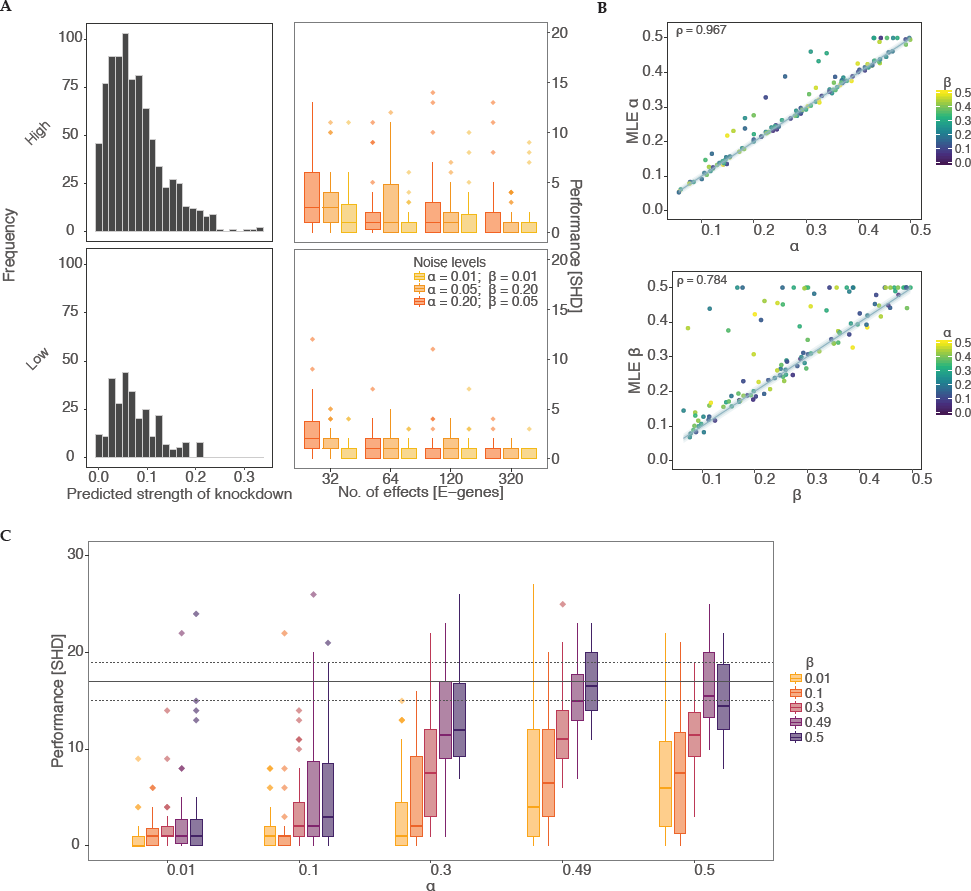
Results from simulation study on DAGs. A. (Left column) Histograms of predicted strength of knockdown depicting the frequency of off-target effects for 30 different DAGs of size *N* = 8, each sampled from *hsa05200* (high density) and *hsa00030* (low density) pathways. (Right column) Structural Hamming distance (SHD) measuring the performance (y-axis) of pc-NEMs on simulated data from the corresponding DAGs with varying number of phenotypic effects (x-axis) and noise levels. The box-plots correspond to the inference from data with 32, 64, 120, and 320 effects and known error rates of *α* = 0.01 and *β* = 0.01 (yellow), *α* = 0.05 and *β* = 0.20 (orange), and *α* = 0.20 and *β* = 0.05 (dark-orange). B. Comparison of 120 different MLE false positive (*α*) and false negative (*β*)rates learned using pc-NEMs against true *α* and *β* used to generate the data. In estimation of *α* (*β*), each point is coloured based on the corresponding *β* (*α*) value used to generate the data. The error rates are inferred from simulated data with 320 phenotypic effects, without *a priori* knowledge of the true network structures. C. Performance (y-axis) of pc-NEMS for network inference without *a priori* knowledge of error rates, on simulated data with 320 phenotypic effects at different noise levels. The horizontal dashed and solid lines correspond to the lower, median, and upper quantile values of SHD for random DAGs.

Finally, to evaluate the robustness of both structure and noise parameter inference, we evaluated the performance of network inference as a function of varying error rates without *a priori* knowledge of the error rates (Fig. 4C). Our method achieved very low values of SHD under extreme values of error rates and continued to consistently perform well up to error rates as high as *α* = 0.5 and *β* = 0.3 or *vice versa* (Fig. S4). This is critical since we observe these levels of noise in the experimental setting and the study confirms that pc-NEMs are robust against a wide range of noise levels.

### 3.3 Application to pathogen infected single-cell data

We applied pc-NEMs to genes associated with *Bartonella henselae* infection. *B. henselae* is transmitted to humans by cat scratches or bites, and causes infections that manifest a broad spectrum of symptoms. Rhomberg et al. (2009) and Truttmann et al. (2011) demonstrated that integrin *β* 1, FAK, Src kinase, paxillin, vinculin, talin1, Rac1, and Cdc42 are required for invasome-mediated uptake of the bacteria. These genes constitute members of the focal adhesion complex and are thus involved in cell attachment and migration. The siRNAs designed to knockdown these genes displayed a high degree of off-targets (off-target frequency = 38), thus making pc-NEMs a good fit for inferring the relations among them.

First, we applied pc-NEMs to the entire data set (Fig. 5A), then we repeated the inference on 100 bootstrap samples of the original data set for robustness assessment (Fig. 5B). All the edges inferred with a threshold of 0.5 on the bootstrap samples with the exception of the edge between paxillin (PXN) and vinculin (VCL) provided support for a subset of the edges inferred in the single run using all the data. The corresponding inferred error rates on the bootstrap samples were *α* = 0.490 ± 0.016 and *β* = 0.211 ± 0.031, respectively. As seen from the simulation studies, these noise levels, although high, are still manageable by the model and the inference algorithm. In this experimental setting, they could be attributed to the lack of coherent biological controls and the pervasively weak signals observed in the Qiagen screens.

**Figure 5:**
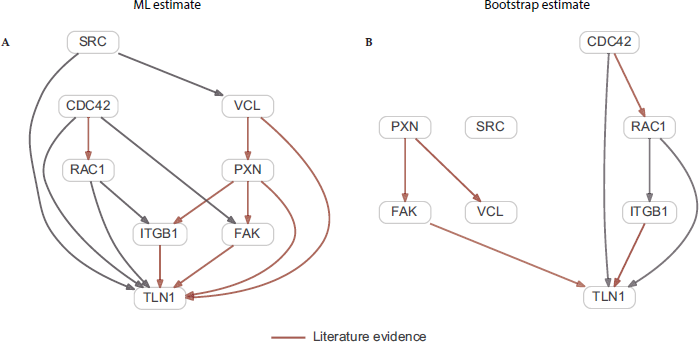
Network inferred using *B. henselae* infection data. Binary gene-level data derived from image-based single-cell data, used for inferring a network of eight genes involved in invasome-mediated uptake of *B. henselae*. A. Maximum likelihood network inferred from entire binary dataset with 288 features resulting from 35 knockdown experiments. B. Network inferred on 100 bootstrap samples of the entire binary dataset. The red arrows correspond to edges with support from literature.

The optimal pc-NEM was able to infer a network in good accordance with knowledge from existing literature. In the inferred network, paxillin is connected to focal adhesion kinase (FAK), vinculin, and talin1 (TLN1); FAK, vinculin and integrin *β*1 (ITGB1) are interacting with talin1. Horton et al. (2015) presented a curated network of the consensus integrin adhesome providing strong evidence for these interactions. Our model also reported an edge between paxillin and integrin *β* 1, a physical interaction suggested by Tanaka et al. (1996).

The regulation of actin cytoskeleton is complex and plays an important role in many cellular processes. pc-NEMs inferred a relation between Cdc42 and Rac1, which is known to be involved in the regulation of the actin cytoskeleton (Verma et al., 2002). Further, our model reported relations between Src kinase (SRC) and vinculin and between FAK and Cdc42. Both these interactions are known to play a role in the context of cellular motility (Ito et al., 1982; Zhang et al., 2004; Zhao et al., 2011).

Some established interactions such as FAK - Src (Mitra et al., 2005) and SRC - integrin *β*1 (Huveneers et al., 2009) were not reported by our model. However, this could be attributed to the low signal to noise ratio as these edges had a bootstrap support of 0.36 and 0.34, respectively.

In addition to the well characterized interactions, our model inferred edges from Cdc42 and Rac1 to talin1, and from Rac1 to integrin *β*1. The PPI network reported by STRING indicates that these associations have been reported in curated databases. Thus, these novel interactions, subject to further experimental investigations, may help provide new insights into the mechanism of *B. henselae* infection.

## 4 Discussion

RNAi screens are highly confounded by off-target effects rendering their analysis challenging. The conventional methods of network inference from perturbation data do not account for these off-target effects, leading to incorrect inference of the network. To address this limitation, here we have developed a novel model based on NEM, called pc-NEM, which handles the combinatorial off-target effects in a probabilistic manner.

In extensive simulation studies, we have demonstrated that by accounting for off-target effects, pc-NEMs improve inference of network topology and error parameters over classical NEMs. We have established a superior performance of pc-NEMs over NEMs in the simulation study over transitively closed networks. While the transitive closure constraint helps explain the nested architecture of the data in the classical setting, it also gives rise to equivalent classes of networks. The probabilistic aspect of the model enables us to relax the strong transitivity constraint of NEMs and still explains the subset relations in the data, thus making the model both biologically relevant and mathematically identifiable. The combinatorial treatment of perturbations empowers us to account for off-target effects, thereby reducing the number of false positive edges.

We have also demonstrated that the model performance is invariant to the density of off-target effects inherent to the data. In contrast to classical NEMs, pc-NEMs can also estimate the error rates in the data as illustrated by the results of our simulation studies. This feature of pc-NEMs is particularly important as it sheds light on the quality of the data or potential model misspecification. Subsequently, we have shown that our model is robust against high noise values. Using pc-NEMs, we reconstructed the network describing the relations between genes involved in *B. henselae* infection from perturbation data with a high degree of off-target effects and high levels of noise. Apart from the well established interactions, we also found novel edges that show potential for improving our understanding of the underlying mechanism. The use of off-target information in combination with the ability to infer error rates reduces the number of false positive edges, making pc-NEMs more suitable for such extreme experimental conditions over classical NEMs.

A challenge for our method lies in scaling with network size. The search space of topologies increases exponentially with the number of signalling genes, thereby requiring longer runs for larger networks. In the current implementation, the computation of marginal likelihood, and the subsequent estimation of effects graph for every sampled topology is expensive. Improving the efficiency of this step is a subject of future research. Furthermore, the model could be extended to handle continuous data sets, enhancing its flexibility.

While the application of the model in the paper is restricted to siRNA off-target effects, it can in general be used in any combinatorial perturbation setting. Especially, in light of double and triple knockdown screens, which are becoming more and more common, pc-NEMs provide a suitable computational framework for network inference.

## Acknowledgements

### Funding

Part of this work has been funded by SystemsX.ch, the Swiss Initiative in Systems Biology, under Grant No. RTD 2013/152 (TargetInfectX – Multi-Pronged Perturbation of Pathogen Infection in Human Cells), evaluated by the Swiss National Science Foundation.

